# Experimental Human Challenge Reveals Distinct Mechanisms of Acquisition or Protection Against Pneumococcal Colonization

**DOI:** 10.1101/459495

**Authors:** Elissavet Nikolaou, Simon P. Jochems, Elena Mitsi, Sherin Pojar, Edessa Negera, Jesús Reiné, Beatriz Carniel, Alessandra Soares-Schanoski, Victoria Connor, Hugh Adler, Seher Raza Zaidi, Caz Hales, Helen Hill, Angela Hyder-Wright, Stephen B. Gordon, Jamie Rylance, Daniela M. Ferreira

## Abstract

Colonization of the upper respiratory tract with *Streptococcus pneumoniae* is the precursor of pneumococcal pneumonia and invasive disease. Following exposure, however, it is unclear which human immune mechanisms determine whether a pathogen will colonize. We used a human challenge model to investigate host-pathogen interactions in the first hours and days following intranasal exposure to *Streptococcus pneumoniae*. Using a novel home sampling method, we measured early immune responses and bacterial density dynamics in the nose and saliva after pneumococcal exposure. We found that nasal colonization can take up to 24 hours to become established. Also, two distinct bacterial clearance profiles were associated with protection: nasal clearers with immediate clearance of bacteria in the nose by the activity of pre-existent mucosal neutrophils and saliva clearers with detectable pneumococcus in saliva at one-hour post challenge and delayed clearance mediated by an inflammatory response and increased neutrophil activity 24 hours post bacterial encounter.

---

The human respiratory tract is a major site of contact with aerosolised bacteria. Acute respiratory tract infections are common, and pneumonia causes more than 1.3 million child deaths annually^1,2^; as well as frequent hospitalisations in at-risk groups such as the elderly, people with chronic lung disease and asthmatics^3^.

The first stage of such infections is the successful colonization of the upper respiratory tract by the pathogen^4^. *Streptococcus pneumoniae* (Spn), the major bacterial cause of pneumonia, inhabits the nasopharynx of 40-95% of infants and 10-25% of adults without causing disease^5^. Colonization is usually asymptomatic in adults but can be associated with mild rhinitis symptoms in children^6^. Different serotypes of pneumococcus may inhabit the nasopharynx at varying densities^7,8^, which is especially common in children^9^. Colonization may continue for a period of weeks or months and be eliminated and reacquired many times during lifetime^10^. As pneumococcal colonization is the primary reservoir for transmission^11^ and a prerequisite of invasive disease^12^, its control is key to preventing disease. Importantly, the determinants of whether exposure to a bacterium leads to colonization have not been identified. Epidemiological studies and pre-clinical studies with animal models point to factors such as host age, immune status, virus co-infection, exposure to antibiotics, smoking and overcrowded living conditions^13,14,15^. However, evidence from *in vivo* human studies is lacking.

Our well-established Experimental Human Pneumococcal Challenge (EHPC) model allows for the rapid, safe and accurate study of bacterial encounter at the nasopharynx in humans^16^. The precise dose and timing of infection are known. Individuals are inoculated with live type 6B pneumococcus and pneumococcal colonization (detection and density), is assessed by nasal washes collected from 48 hours onwards post-exposure. After this point, approximately 50% of individuals are identified as carriers, but the early kinetics of bacterial clearance or colonization onset have not been assessed. Here, healthy adult volunteers were challenged with *S. pneumoniae*, after which they self-collected saliva and nasal lining fluid (nasosorption) samples for bacterial kinetics and immune monitoring in the first 48 hours. We compared non-colonized and colonized subjects to identify potential correlates of protection and to define the kinetics of colonization. Our hypothesis was that mucosal innate responses would be predictive of protection against colonization and the movement of bacteria from the nose (site of exposure) to the saliva within the first 48 hours would be associated with less effective immunity and potentially with colonization.

## Results

### Home sampling method can be used to collect samples at defined timepoints

Forty volunteers aged 18–32 years were screened for pre-existing colonization with pneumococcus and non-colonized participants were inoculated with 6B pneumococcus, as previously described^16^. Colonization status was defined by classical microbiology culture of *S. pneumoniae* serotype 6B in nasal wash samples collected at day 2, 6, 9 and 14 post exposure. Volunteers with negative samples at all time points were defined as culture-negative and those with a positive sample at any time point were classified as culture-positive. Following exposure, 30 volunteers remained culture-negative (75%) and 10 became culture-positive (25%) (Table S1). Colonization status was confirmed by *lytA* qPCR in nasal wash.

Saliva and nasal lining fluid samples were obtained before exposure (Time=0 hours, baseline). Volunteers collected their own saliva into pre-prepared tubes at 1, 2, 4, 8, 12, 24, 36, and 48 hours, and anterior nasal lining fluid (by nasosorption strip)^17^ at 24 and 48 hours. A subgroup of 33 volunteers self-collected additional nasal lining fluid samples at 4 and 8 hours (Figure 1A). To monitor compliance, volunteers were instructed to record sample collection times using their mobile phones, sending pictures of the collected sample to the research team at the time of sample collection. Samples from two volunteers were excluded (5%) due to poor compliance detected by temperature monitoring (half of their samples were stored at room temperature, Figure S1A). Therefore, data from 38 volunteers were evaluated in this study (Figure S1B); 29 culture-negative and 9 culture-positive volunteers (Table S1). This high rate of compliance indicates the successful application of our novel sampling method.

**Figure 1:**
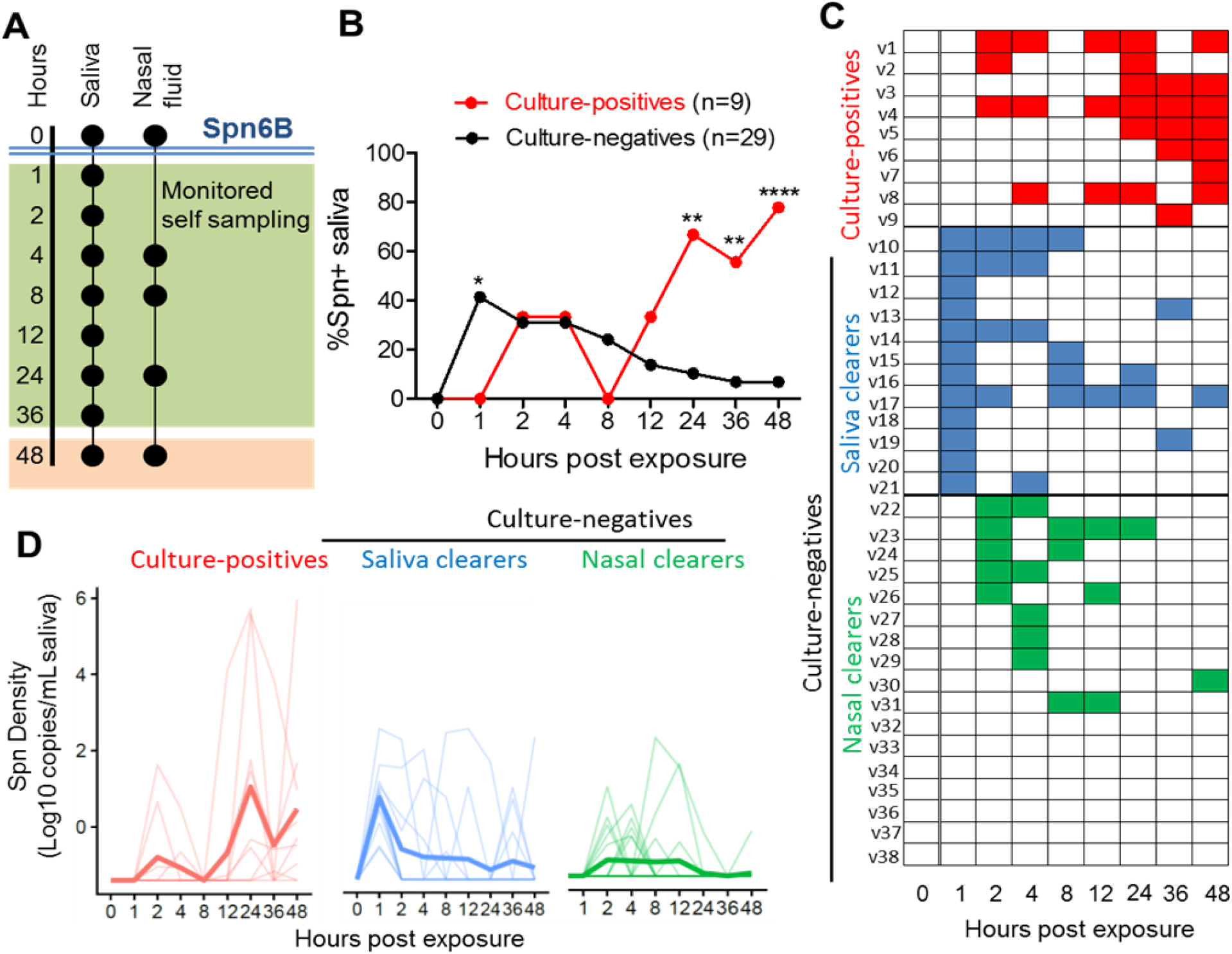
Kinetics of early pneumococcal detection in saliva. (A) Sample collection set-up. Saliva and anterior nasal lining fluid samples were collected before inoculation T=0 hours (baseline) in clinic. Volunteers were asked to self-collect saliva at 1, 2, 4, 8, 12, 24, 36, 48 hours and anterior nasal lining fluid at 4, 8, 24, 48 hours post-inoculation. **(B) Frequency of Spn6B pneumococcus in saliva.** Detection of pneumococcal DNA in saliva was determined by Spn6A/B qPCR. Nine culture-positive and 29 culture-negative volunteers were assessed. The number of volunteers with Spn6B hit (CT <40) in each timepoint is expressed as a percentage (%) of the total number of volunteers. Statistical significance based on Fisher’s exact test *P<0.05, **P<0.001 and ****P<0.00001 **(C) Heatmap showing individual saliva profiles.** Three distinct saliva profiles were defined. Culture-positive, red; volunteers who were identified to be experimentally colonized with pneumococcus at Day 2 or later using classical microbiology (n=9), Saliva Clearers, blue; volunteers with detectable by qPCR Spn levels at 1 hour after exposure (n=12) and Nasal Clearers, green; Volunteers with detectable Spn levels by qPCR later than 1 hour after exposure (n=17). **(D) Density levels of pneumococcal Spn6A/B PCR in saliva.** Spn6B density was expressed as DNA copies per volume (mL) of saliva. If no Spn was detected, the density was set as 0 CFU/mL. Data was log transformed after adding 1 to all values to allow transforming 0 values. Individual volunteers and the mean of log-transformed values are shown.

### Raw and culture-enriched DNA extraction in saliva are complementary for Spn detection in saliva

To detect Spn presence, pneumococcal genomic DNA was extracted from both raw and culture-enriched (blood agar plate with gentamicin) saliva samples, as the later has been shown to be the best method for pneumococcal detection in saliva^18^. In total, 58 Spn hits were detected using a Spn 6A/B capsule-specific qPCR (26 in raw samples, 16 in culture-enriched, and 16 in both, Figure S1C). Spn density from culture-enriched samples was significantly higher (lower Ct values) than from raw samples (Figure S1D). Consequently, samples with a positive result using either method were included in further analyses.

### Two distinct profiles of bacterial clearance kinetics associate with protection against colonization

To investigate Spn clearance kinetics, we assessed the presence of pneumococcus DNA in saliva in the first 48 hours after exposure by qPCR (Figure 1A). At one hour, amongst culture-negative volunteers, Spn DNA was detected in the saliva of 12/29 (41.4%) volunteers (Figures 1B and 1C). This suggests that two distinct profiles of protection against colonization exist. We named these two groups “nasal clearers” (individuals with no detectable Spn DNA in 1-hour post-exposure saliva samples) and “saliva clearers” (individuals with detectable Spn DNA in 1-hour post-exposure saliva samples). Nasal clearers showed similar levels of Spn density between 2 and 12 hours (Figure 1D). Saliva clearers showed a peak of Spn density (DNA copies/ml saliva) at 1 hour after exposure and followed by gradually decrease of Spn density after 1 hour.

### Establishment of pneumococcal colonization takes up to 24 hours

Amongst those subsequently culture-positive volunteers, Spn DNA was not detected in any 1hr saliva samples (Figures 1B and 1C). At 24 hours, Spn DNA was detected in 6/9 (67%) of culture-positive volunteers, but in only 3/29 (10%) of culture-negative volunteers (Figures 1B and 1C). Moreover, at 48 hours, Spn DNA was detectable in the saliva of most culture-positive 7/9 (77.8%), but few culture-negative 2/29 (6.9%) volunteers (Figure 1B). The mean density started increasing significantly in culture-positive volunteers only from 24 hours. This suggests a gradual colonization process that takes at least 24 hours.

### Bacterial DNA detection in the nose

We then evaluated the presence of pneumococcal DNA in the nasal lining fluid during the first 48 hours post exposure by qPCR, as described above (Figure 1A). Volunteers missing one or more home samples (n=12; 5 culture-positive and 7 culture-negative volunteers) were excluded from this analysis. At 4 hours post exposure, Spn DNA was detected in the nose of 20/21 (95%) culture-negative, and in all culture-positive 5/5 (100%) volunteers. At 8 hours, Spn DNA was detected in 10/11 (91%) nasal clearers, 6/10 (60%) saliva clearers, and 5/5 (100%) culture-positive volunteers (Figure 2A). Thus, in 40% of saliva clearers, Spn DNA was completely removed from the exposure site (the nose) within 8 hours. At 24 and 48 hours Spn DNA was detected in nasal lining fluid from only 1/5 culture-positive volunteers, indicating that in 4/5 volunteers the bacteria had either migrated posteriorly, or become attached or internalised at the epithelium. For saliva and nasal clearers, Spn DNA was detected in the nose of 5/10 (50%) and 5/11 (45%) volunteers at 24 hours and 2/10 (20%) and 2/11 (18%) volunteers at 48 hours post exposure respectively. There was no difference in late nasal clearance profile between the two groups (Figure 2B).

**Figure 2:**
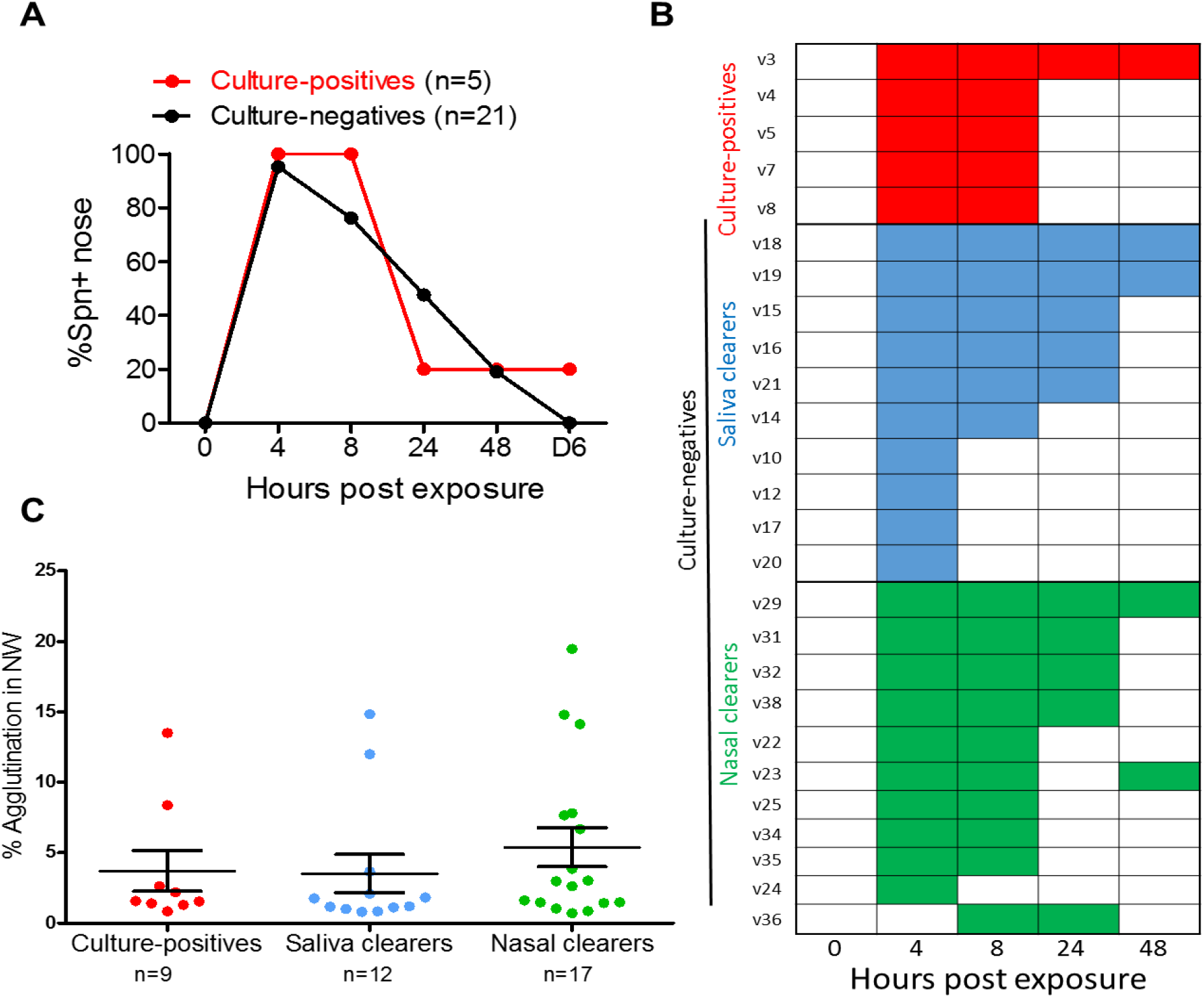
Kinetics of early pneumococcal detection in anterior nasal fluid. (A) Frequency of Spn presence in nasal fluid after challenge.
 Nasal fluid samples were collected before (T0) pneumococcal exposure and at 4, 8, 24 and 48 hours post exposure. Detection of pneumococcal DNA in nasal fluid was determined by Spn6A/B qPCR. Volunteers with a missing sample were excluded from this analysis. Therefore, 5 culture-positive and 21 culture-negative volunteers were assessed. The number of volunteers with a Spn6B hit in each timepoint expressed as a percentage (%) of the total number of volunteers. **(B) Heatmap showing individual nasal profiles distributed in three distinct nasal profiles.** 5 culture-positives, 10 saliva clearers and 11 nasal clearers. Each detected hit was colour coded; red for culture-positives, blue for saliva clearers and green for nasal clearers. **(C) Levels of agglutination capacity in nasal wash at baseline. Each dot represents one volunteer.** Mean ± SEM are shown for each of the 3 groups.

### Pneumococcal agglutination was not associated with early pneumococcal profiles

Agglutination of pneumococcus by polysaccharide 6B (PS6B) specific IgG antibodies has been previously shown to protect against 6B pneumococcus colonization in the context of pneumococcal conjugate vaccination^19^. Also, mucus is known to be able to trap pathogens^20^ and to bind pneumococcus through carbohydrate motifs^21^. To investigate whether agglutination correlates with protection or colonization clearance profiles, agglutination capacity was measured in baseline nasal wash samples and compared with the levels of mucin (MUC5AC) and Spn polysaccharide specific IgG antibodies (PS6B) in these samples. Baseline nasal wash agglutination capacity was correlated significantly with high levels of MUC5AC (Figure S2A and S2B), but not with PS6B-specific IgG levels (Figure S2C and S2D).The agglutination capacity (Figure 2C), MUC5AC, and PS6B-specific IgG levels were not siginficantly different between the three groups in nasal wash at baseline.

### Neutrophil activity contributes to protection against establishment of colonization in nasal clearers

Neutrophils are abundantly present in the adult human nose even in the absence of pneumococcal colonization^17^. Neutrophils are activated by bacterial encounter, releasing myeloperoxidase (MPO) during degranulation^22^. To investigate if neutrophil activity at the time of bacterial encounter contributes towards pneumococcal clearance, the abundance of neutrophils and MPO levels were measured at baseline. Nasal immune and epithelial cells were measured from nasal curettes (Figure S3A)^17,23^. The absolute number of activated neutrophils were identified by measuring CD66b^Hi^ granulocyte levels, a marker for neutrophil activation^24^. We also analysed total number of granulocytes and the expression levels of CD66b within the granulocyte population based on their MFI (Mean Fluorescence Intensity) (Figure S3A).

Nasal clearers showed at baseline 10-fold and 11-fold higher levels of activated (CD66b^Hi^) and total granulocytes, respectively, compare to culture-positive (Figure 3A and Fig S3B) volunteers. Moreover, the granulocytes of nasal clearers had increased expression of CD66b compared to saliva clearers (Figure S3C). This finding was supported by increased MPO levels at baseline in nasal wash in nasal clearers compared to culture-positives and saliva clearers (Figure 3B). There was a significant correlation between numbers of CD66b^Hi^ granulocytes and MPO levels in nasal wash (Figure 3C). There was no significant difference in levels of any other measured cell type (B cells, T cells, epithelial cells and monocytes) between the three groups (Figures S3D-G) indicating that increased levels of neutrophils at baseline are protective against pneumococcal colonization in nasal clearers.

**Figure 3:**
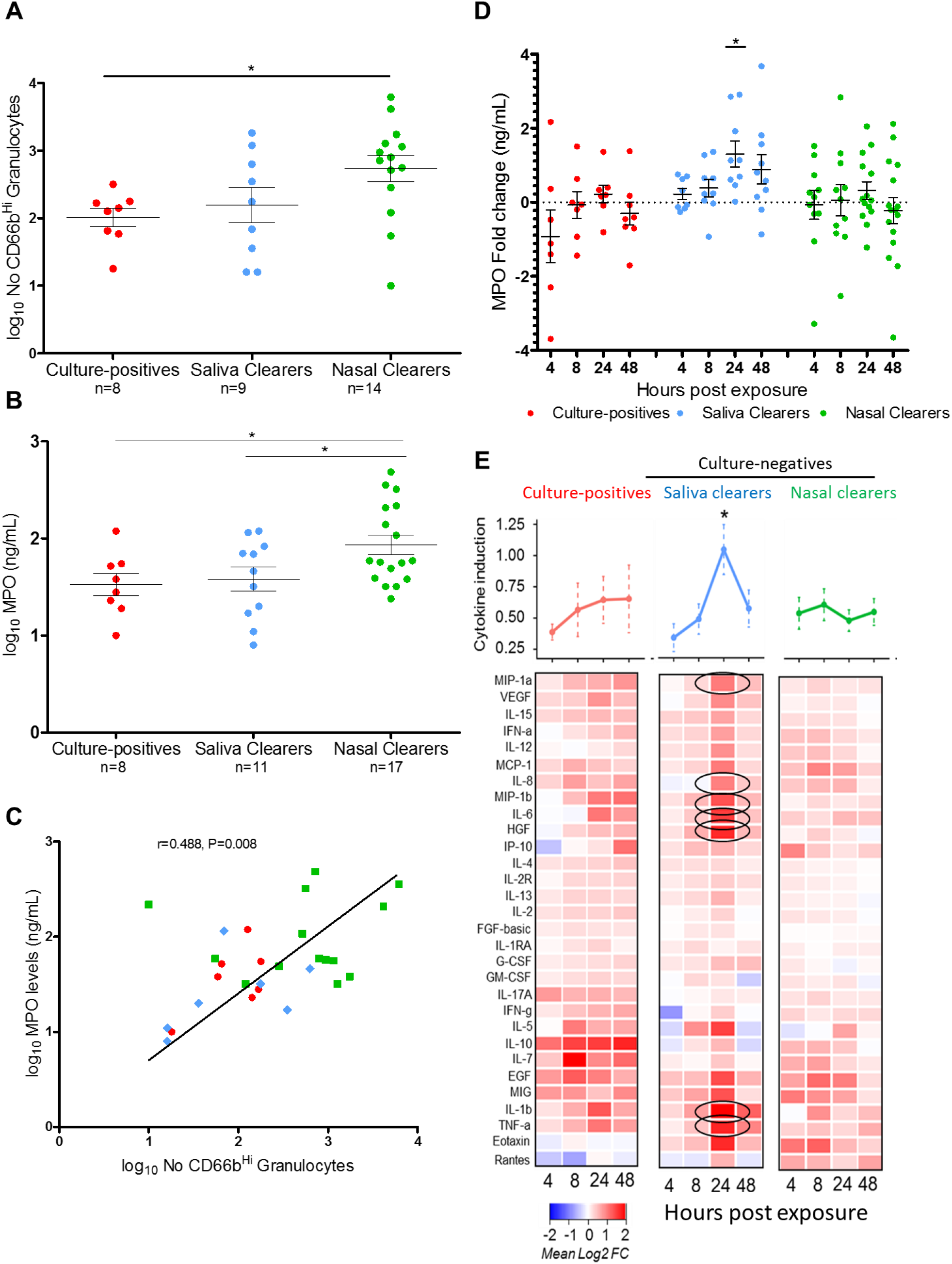
Association of mucosal immune factors with early pneumococcal colonization profiles. (A) Neutrophil number in nasal scrapes prior to pneumococcal exposure. Abundance of CD66b-high cells (activated granulocytes) at baseline were measured by flow cytometry. Eight culture-positives, 9 saliva clearers and 14 nasal clearers were assessed (*P=0.020, unpaired t test). Data was log transformed and individual volunteers and mean ± SEM are represented. **(B) MPO levels in nasal wash prior to pneumococcal exposure.** MPO levels at baseline were measured by ELISA. Eight culture-positives, 11 saliva clearers and 20 nasal clearers were assessed. MPO levels in nasal wash were significantly increased in nasal clearers compared to culture-positives (* P=0.023, unpaired t test) as well as saliva clearers (* P=0.036, unpaired t test). Data was log transformed and individual volunteers and mean ± SEM are represented.) **(C) Correlation between MPO levels in nasal wash and number of activated granulocytes (CD66b^Hi^) in nasal scrapes prior to Spn challenge.** Data was log transformed. A Pearson test was used (r=0.488, **P=0.008). Culture-positives; red circles, saliva clearers; blue rectangular and nasal clearers; green squares. **(D) MPO levels in nasosorption during the first 48 hours.** Myeloperoxidase levels in nasal fluid were measured by ELISA. Fold change of MPO levels was calculated for each group according to each group’s baseline MPO levels. Data was log transformed and are represented as mean ± SEM. In saliva clearers, levels of MPO rose after exposure, peaking at 24hours after pneumococcal exposure (*P=0.013, unpaired test to baseline). **(E) Heatmap showing cytokine induction in nasal lining fluid following Spn challenge.** Abundance of 30 cytokines was measured by Luminex at 4, 8, 24 and 48 hours after pneumococcal exposure and normalized to baseline levels for each subject. The mean of log2-tranformed fold-changes are shown per timepoint for each of the three groups. Statistical analysis was performed using R Studio. Statistical significance based unpaired t test comparing to baseline, followed by multiple testing correction (Bejamini-Hochberg), with significance P ≤ 0.05, with 7 cytokines significantly upregulated at 24 hours (black circles); IL-6, IL-1β, MIP-1α/CCL3, MIP-1β/CCL4, HGF, TNF-α and IL-8. Saliva clearers showed a significant total induction score of cytokines at 24 hours post exposure compared to nasal clearers and culture-positives (*P=0.027, one-way ANOVA).

To assess early neutrophil responses following exposure, we measured levels of MPO in nasal lining fluid in the first 48 hours post exposure (Figure 3D). All three groups had comparable levels at baseline. Levels of MPO remained the same during the first two days in nasal clearers and culture-positive volunteers. In saliva clearers, however, levels of MPO rose after exposure and reached a significant peak at 24hours, 3-fold higher than at baseline. Thus, an induced neutrophil response following exposure supports the role of this cell population in protection against delayed colonization in saliva clearers.

### A robust inflammatory response is associated with protection from colonization in saliva clearers

To investigate whether a prompt immune response contributes towards protection from colonization by pneumococcus, we measured the abundance of 30 cytokines in nasal fluid using Luminex. A total cytokine induction score was calculated based on all cytokines similar to that previously described for gene expression^25^. Distinct cytokine profiles were observed within the three groups (Figure 3E). Culture-positive volunteers had no significant cytokine induction at any time point measured during the first 48 hours, analogous to prior findings at later timepoints^26^. Similar to culture-positive volunteers, nasal clearers showed no significant induction of cytokines at any time point. In contrast, saliva clearers showed a significant induction of cytokines at 24 hours post exposure, with 7 cytokines (IL-6, IL-1β, MIP-1α/CCL3, MIP-1β/CCL4, HGF, TNF-α and IL-8) significantly upregulated (Figure 3E). The induction of all of these cytokines except IL-6 strongly correlated at 24 hours with MPO production in nasal lining fluid (Figure S3H). These results suggest that saliva clearers induce a strong pro-inflammatory response, which associates with early neutrophil responses, leading to protection against colonization.

## Discussion

To develop effective protective interventions against disease, it is vital to understand which mechanisms play a crucial role in host-pathogen interactions by protecting against establishment of colonization following bacterial encounter. Here we investigated human-pneumococcus interactions in the first 48 hours following controlled intranasal exposure. We tested the hypothesis that volunteers who are susceptible to colonization have a distinct profile of Spn DNA kinetics in the nose and saliva from the culture-negative group.

To our surprise we observed 3 distinct groups: culture-positive volunteers with a high concentration of Spn DNA in the nose 24 hours following nasal exposure, and culture-negative volunteers, among whom there were two groups – one who had detectable Spn DNA in the saliva 1 hour after exposure and another group who did not. We named these two culture-negative groups saliva clearers and nasal clearers, respectively.

Culture-positive volunteers showed no fast initial clearance (appearance in saliva) as well as a lack of cytokine and neutrophil responses during the first hours and days following exposure. Colonization was therefore established in that period. We then observed pneumococcal density diminishing in the anterior nose (exposure site) by 8 hours suggesting that migration towards the nasopharyngeal site may take place by this time point. Alternatively, the absence of pneumococcus from the anterior nose may reflect strong epithelium binding or even internalization of the bacteria. Indeed, pneumococcus was found to attach to epithelium of the inferior nasal turbinate in the EHPC model during colonization^27^. Although we see differences these are small numbers. In this group, colonization takes at least 24 hours to be fully established, at which point Spn can be robustly measured in saliva.

Nasal clearers showed a strong baseline neutrophil activation. These volunteers did not develop a significant pro-inflammatory response or had bacterial movement to the saliva within one hour. It is possible that the high level of baseline activated nasal neutrophils prevent sensing by epithelia or other cell types such as monocytes as well as the movement to the saliva. Thus, cytokine responses were not induced in these volunteers. This indicates that neutrophils play a key role in the early control of human pneumococcal colonization. This has not been appreciated in murine models of infections, as neutrophils are present in the nasal lumen of humans^28^, whereas the nasal mucosa of naive mice has no neutrophils^29^ and the cell influx occurs only following bacterial challenge. Indeed, the relative prevalence of pneumococcal serotypes in causing colonization in the community associates with their relative capacities to resist neutrophil-mediated killing *in vitro*^30^.

Saliva clearers demonstrated an initial fast movement of the bacteria to the saliva, which we postulate is due an effective nasal mucociliary activity. In a subset of these (40%), Spn was completely cleared from the nose within 8 hours but it could still be detected in all others at 24 hours. Moreover, saliva clearers induced a strong pro-inflammatory response in the first day post exposure, which we were able to measure at 24 hours. This is associated with a concurrent induction of neutrophil responses. Taken together, this suggests that this group is protected likely by mucociliary clearance which is then supported by a strong neutrophil response to pneumococcus.

Mucus components can influence the bahavior of pathogenic bacteria by increasing their virulence expression, adhesion, motility, proliferation or growth^31^. While bacterial movement suggests that mucociliary clearance could be involved in protection of a subset of volunteers, we were not able to establish a direct link between pneumococcal agglutination and early pneumococcal clearance by mucocilliary movement. The observation that MUC5AC levels correlated with agglutination capacity supports the role for mucus in binding pneumococcus^30^. Neither agglutination nor agglutinating factors (mucus, PS6B), however, associated with early pneumococcal clearance profiles in these unvaccinated volunteers. This result stands in contrast to a previously demonstrated effect of agglutinating polysaccharide specific IgG and its protective role in pneumococcal colonization in the context of vaccination^19^: in this case the levels of polysaccharide antibody are very highly increased at which point mucocilliary clearance facilitated by agglutination may play a role in preventing pneumococcal colonization. The latter indicates that these results might be relevant for vaccine-induced rather than naturally acquired immunity. Alternatively, mucociliary clearance may be important in this defence process. Rapid bacterial migration was protective in the absence of agglutination capacity, nasal ciliary function was not measured in this study. Smokers^32^ and individuals with respiratory tract viral infections, particularly respiratory syncytial viral^33^ have decreased mucociliary beating and are more prone to pneumococcal colonization.

Our data describe a novel home sampling method for defining the early events after experimental pneumococcal challenge. Stable colonization following pneumococcal encounter takes at least 24 hours. We demonstrated that there are distinct profiles of pneumococcal colonization kinetics, distinguished by speed of clearance, local phagocytic function, and acute mucosal inflammatory responses which may either recruit or activate neutrophils. These results highlight that when establishing correlates of protection against bacterial colonization, which can be used to inform design and testing of novel vaccine candidates, multiple mechanisms of protection must be considered.

## Methods

### Experimental Human Pneumococcal Colonization Study - Recruitment of volunteers and ethical statements

Volunteers were enrolled from a concurrent clinical trial conducted in 2016-2017. Details on participant demographics and study’s design have been previously described^34^. Limited subject characteristics for this cohort are shown in Table S1. Briefly, volunteers were screened for *S*. *pneumoniae* colonization (natural carriers) and after 6 days were intranasally inoculated with Spn6B serotype (strain BHN418, GenBank ASHP00000000.1) (8×10^4^CFU/100ul per nostril). Colonization was assessed by classical microbiology culture in nasal washes collected at 2, 6, 9, 14, 21 and 27 days post exposure. Pneumococcus type was confirmed by latex agglutination (Statens Serum Institute, Copenhagen, Denmark).

Ethical approval was given by local NHS Research and Ethics Committee (REC) (14/NW/1460) and the study was registered on the European Clinical Trials Database (EudraCT Number 2014- 004634-26). All experiments were conformed to the relevant regulatory standards (Human Tissue Act, 2004). Informed consent was obtained from all volunteers.

### Home sampling procedure

Volunteers were given written instruction sheets for sample self-collection and picture taking/sending via mobile messaging application, plus a sample data collection form (time planner). Volunteers were given a transport bag containing 8 × 10mL saliva collection tubes and funnels (plus one spare) (Isoxelix^TM^, Cell Projects Ltd, Kent, UK) with 1mL skim milk, tryptone, glucose, and glycerine (STGG) media with 50% glycerol for bacterial preservation and transport, 4 adsorptive matrix filter strips (Nasosorption^TM^, Hunt Developments Ltd, West Sussex, UK) for nasal fluid collection, 2 ice packs for keeping samples cooled during the day, 1 plastic box for sample storage and a USB temperature data logger thermometer (Woodley Equipment Company Ltd., Lancashire, UK) for temperature monitoring. For nasal lining fluid collection, the matrix strip was inserted into the nostril and held against the nasal lining for 2 min and then placed in its transport tube. For saliva collection, volunteers spat 1 mL (maximum) of saliva into the STGG tube. Both sample collection methods were demonstrated to the volunteers at baseline sample collection). Optimization experiments were performed to investigate optimal storage of the samples by spiking saliva with different Spn concentrations. Results indicated that Spn density was lower to samples stored at ambient temperature, however similar Spn density was detected to samples stored either in fridge or freezer. Samples taken were stored in the volunteer’s home freezer overnight and transported to the lab at their day 2 clinic visit, where they were immediately stored at -80°C until use.

### Bacterial DNA extraction from saliva samples

Bacterial genomic DNA was extracted from raw and culture-enriched saliva samples. On the day of the extraction, saliva samples were thawed for 30min in RT and vigorously vortexed for 20sec. 200μl of raw saliva was used for DNA extraction. In addition, for culture-enrichment step, 10μl of raw saliva was diluted with 90μl of saline and cultured on Columbia blood agar supplemented with 5% horse blood (PB0122A, Oxoid/Thermo Scientific) and 80μl gentamycin 1mg/mL (G1264-250mg, Sigma-Aldrich co Ltd). Plates were incubated overnight at 37°Cand 5% CO_2_. The remaining raw saliva was stored at -80°C. After incubation, all bacterial growth was harvested into 2mL STGG and vigorously vortexed until homogenise. 200μl of culture-enriched saliva was used for DNA extraction and the remaining samples were stored at -80°C.

For both raw and culture-enriched saliva samples, thawed suspensions were pelleted in a 1.5mL tube at 20,238xg for 10 min. The pellet was resuspended in 300μl of lysis buffer with protease (Agowa Mag mini DNA extraction kit; LGC Genomics, Berlin, Germany), 100μl of sterilized zirconia/silica beads (diameter of 0.1 mm; Biospec Products, Bartlesville, OK, USA), and 300μl of phenol (Phenol BioUltra; Sigma-Aldrich, Zwijndrecht, The Netherlands). The sample was mechanically disrupted by bead beating in a TissueLyser LT (Qiagen, Venlo, The Netherlands) twice at 50 Hz for 3min. After 10 min centrifugation at 9,391g, the aqueous phase was transferred to a sterile 1.5mL tube. Binding buffer was added at twice the volume of the aqueous phase plus 10μL of magnetic beads, after which the sample was incubated in a mixing machine (^~^265 rpm) for 30min at RT. The magnetic beads were washed with 200μL of both wash buffer 1 and wash buffer 2, and eluted with 63μL of elution buffer, according to the manufacturer’s instructions. For optimization experiments, DNA from all raw saliva samples was also extracted using QIAamp DNA Mini Kit (Qiagen, Manchester, UK), following manufacturer’s instructions. This method showed same results with the one described previously in this section.

### Bacterial DNA extraction from nasal fluid pellet

On the day of the extraction, nasosorption filter strips were thawed for 30min in RT. 100μl of assay diluent added to the filter and centrifuged at 1,503g for 10min. After centrifugation the eluted liquid was moved to a clean Eppendorf tube and centrifuged at 16,000g for 10 minutes at 4°C. The supernatant was removed and used for cytokine analysis, whereas the pellet was used for DNA extraction. Bacterial genomic DNA was extracted from the nasal fluid pellets using the same method as described above (saliva samples).

### Quantification of pneumococcal DNA by qPCR in saliva and nasal fluid pellet samples

Colonization density was determined by 6A/B-specific qPCR targeting the *CpsA* gene (Azzari et al., 2010) using the Mx3005P system (Agilent Technologies, Cheadle, UK). The primers and probe sequences were as follows: forward primer, 5’-AAGTTTGCACTAGAGTATGGGAAGGT-3’; reverse primer, 5’-ACATTATGTCCATGTCTTCGATACAAG-3’; probe, 5’-(FAM)-TGTTCTGCCCTGAGCAACTGG-(BHQ1)-3’^29^. The 25μL PCR mix consisted of 12.5μL 1 × TaqMan Universal PCR Master Mix (Life Technologies Ltd, Paisley, UK), 0.1μL *100μ*M each primer, 0.05μL 100μM probe, 9.75μL molecular graded water (Fisher Scientific, Loughborough, UK) and 2.5μL of the extracted DNA. Thermal cycling conditions were: 10min at 95°Cand 40 cycles of 15secs at 95°Cand 1min at 60°C. A negative DNA extraction control (parallel extraction from sample buffer only), a qPCR negative control (master mix only) and three extractions of each sample were amplified. A standard curve of a ten-fold dilution series of genomic DNA extracted from Spn6B was used. The genomic DNA was extracted with the QIAamp DNA Mini Kit (Qiagen, Manchester, UK) and quantified with a spectrophotometer (Nanodrop ND-1000; Thermo Fisher Scientific, Landsmeer, The Netherlands). To convert the weight of pneumococcal DNA to number of *S. pneumoniae* DNA copies, the weight of one genome copy of TIGR4 was used to calculate the genome length in base pairs times the weight of a DNA base pair (650 Da). Samples were considered positive if two or all triplicates yielded a C_T_ < 40 cycles. Multiple experiment analysis was performed, and cross experiment threshold was calculated by using inter-run calibrators.

### Human Myeloperoxidase ELISA

Levels of myeloperoxidase were determined using the Human Myeloperoxidase DuoSet ELISA Kit (R&D Systems, Abingdon, UK). 96-well ELISA plates were coated with 4μg/ml capture antibody in PBS at RT overnight. Plates were washed 3 times with PBS (Sigma-Aldrich Co Ltd, Irvine, UK) containing 0.05% Tween-20 (Sigma-Aldrich Co Ltd, Irvine, UK) between each step. Wells were blocked with 1% BSA in PBS for 1 hour at RT. Samples and standards were diluted in 1% BSA-PBS in the pre-coated plates and incubated at RT for 2 hours. Detection was performed by incubating plates with detection antibody at 50ng/ml for 2hours at RT, followed by 20min incubation with Streptavidin-HRP (1:200) (Fisher Scientific, Loughborough, UK) at RT. Signal was developed using TMB-Turbo Substrate (Fisher Scientific, Loughborough, UK) for 20min and stopped by adding 2N H_2_SO_4_ in a 1:1 ratio. Optical density reading was performed at 450nm and corrected for optical imperfection (540nm). All samples were run in duplicate. Results are expressed as μg/ml and calculated using MPO standard curve.

### Agglutination assay by flow cytometry

Spn6B cells were grown to mid-log-phase and stored at −80°C in glycerol until use as described before^19^. For agglutination assays with human nasal wash samples, cells were thawed and washed with PBS and 2μl of bacteria was incubated with 48μl of concentrated nasal wash supernatant (1ml of nasal wash concentrated to 50μl using vacuum concentrator RVC2-18) and dialysed overnight in PBS using Slide-A-Lyser Dialysis Units (TermoScientific). Antiserum to group 6 (Statens Serum Institute, Neufeld antisera to group 6) was used as a positive control and Anti-Hep-A purified human IgG was used as a negative control (using sepharose and pooled sera from HepA vaccinated volunteers). Samples were vortexed lightly and incubated for 1.5h at 37°C, 5%CO_2_.

Cells were fixed with paraformaldehyde (PFA) and analysed on a BD LSR II Flow Cytometer (BD Biosciences, San Jose, CA, USA). Bacterial population was gated in the Forward scatter (FSC) and Sideward scatter (SSC) dot plot referring to cell size and granularity. PMT voltages and threshold were gated on negative control bacteria. 30,000 events of each sample were measured by triplicate using FacsDiva Software 6.1 (BD Biosciences, San Jose, CA, USA). Agglutination was quantified by calculating the proportion of the bacterial population with altered FSC and SSC and values were expressed as % of agglutination, as previously described^35^. All samples were analysed in duplicate and 30,000 events were acquired using FacsDiva Software 6.1 (BD Biosciences, San Jose, CA, USA). Analysis was performed using FlowJo software version 10.0 (Tree Star Inc, San Carlos, CA, USA).

### Nasal cells processing and flow cytometry

Cells were dislodged from the curette by repeated pipetting with PBS+ as described previously^17^. Cells were spun down at 440g for 5 min and resuspended in PBS++ containing LIVE/DEAD^®^ Fixable Aqua Dead Cell Stain (ThermoFisher). After 15 min incubation on ice, an antibody cocktail which included Epcam-PE, HLADR-PECy7, CD66b-FITC, CD19-BV650 (all Biolegend), CD3-APCCy7, CD14-PercpCy5.5 (BD Biosciences) and CD45-PACOrange (ThermoFisher) was added to the cells. Following a further 15 min incubation on ice, cells were filtered over a 70μm filter (ThermoFisher). Cells were spun down (440g for 5 min), resuspended in PBS++ and acquired on a flow cytometer (LSRII, BD). Samples with less than 500 immune cells or 250 epithelial cells were excluded from further analysis. Flow cytometry data was analysed using Flowjo V. 10 (Tree Star Inc, San Carlos, CA, USA).

### Cytokine Analysis

Nasal washes were centrifuged at 1,503g for 10 mins and the extracted supernatant was stored at -80°Cuntil use. Human MPO was measured according to manufacturer’s instructions. The human magnetic 30-plex cytokine kit (ThermoFisher) was used to detect thirty cytokines simultaneously on a LX200 with xPonent3.1 software (Luminex) following manufacturer’s instructions. Cytokines for samples with a CV >25% were excluded from further analyses.

### Heat map generation and total cytokine score

Heat map representations were generated using R. Fold change concentrations to baseline were calculated for each individual and log_2_-transformed. An average fold change for each group for each timepoint was then calculated. A total cytokine score was calculated as previously described^35^. In brief, the average log_2_ fold change of all upregulated cytokines for a given individual was calculated, by replacing downregulated cytokines with a value of 0 and then taking the average cytokine log_2_ fold change per sample.

### Quantification and Statistical analysis

Statistical analysis was performed by GraphPad Prism version 5.0 (California, USA) and R software. Data was log transformed where appropriate. To distinguish between parametric and non-parametric data a Kolmogorov-Smirnoff test was performed. If two parametric groups were compared, a two-tailed *t* test was used for unpaired and paired groups. If two non-parametric groups were compared, a Mann-Whitney or Wilcoxon test was used for unpaired and paired groups respectively. If multiple unmatched groups were compared, a one-way ANOVA (followed by a Tukey’s post-test) or Kruskal-Wallis test (followed by a Dunn’s post-test) was used for parametric or non-parametric groups respectively. For Luminex data, a Benjamini-Hochberg correction was used, to account for testing of 30 cytokines simultaneously. To quantify association between groups, Pearson or Spearman correlation test was used for parametric or non-parametric groups, respectively. Differences were considered significant if p ≤ 0.05.

## Acknowledgments

We would like to thank all the volunteers for their participation and very good compliance with study’s procedure. Also, Catherine Lowe and Rachel Robinson for helping collecting home samples, Prof Adam Finn for helpful discussions on the validation of home sampling methods and Dr Alison Isaacs for helpful review of the manuscript. Flow cytometric acquisition was performed on a BD LSR II funded by a Welcome Trust Multi-User Equipment Grant (104936/Z/14/Z). This work was supported by the Medical Research Council (grant MR/M011569/1) and Bill and Melinda Gates Foundation (grant OPP1117728). The funders had no role in study design, data collection and analysis, decision to publish, or preparation of the manuscript.

## Author Contributions

E.N., S.P.J., E.M., S.P. and D.M.F. designed the experiments and wrote the study protocols. S.P.J and D.M.F. did the initial investigation for the home sampling method. E.N and S.P.J. supervised the clinical study and laboratory work. E.N., S.P.J., E.M., S.P., E.Ne., J.Re., B.C., A.S.S, V.C., H.A., C.H., H.H., S.R.Z. and A.H.W participated in site work including laboratory processing, data collection and challenge preparation. E.N., SP.J., E.M. and S.P. performed statistical analyses. E.N. wrote the original manuscript. E.N., S.P.J., E.M., S.P., E.Ne., J.Re., B.C., A.S.S, S.B.G, V.C., H.A., C.H., H.H., S.R.Z., A.H.W., S.B.G, J.R. and D.M.F reviewed and edited the manuscript. All authors significantly contributed to interpretation of the results, critically revised the manuscript for important intellectual content and approved the final manuscript.

## Tables with titles and legends

**Table S1.** Volunteers’ demographic data.

| **Volunteer No** | **Gender** | **Age (years)** | **Colonization Status**<br>**(Culture)** |
| --- | --- | --- | --- |
| 1 | F | 19 | Positive |
| 2 | M | 20 | Positive |
| 3 | F | 21 | Positive |
| 4 | M | 20 | Positive |
| 5 | F | 20 | Positive |
| 6 | M | 20 | Positive |
| 7 | M | 19 | Positive |
| 8 | F | 22 | Positive |
| 9 | F | 22 | Positive |
| 10 | F | 20 | Negative |
| 11 | F | 22 | Negative |
| 12 | F | 22 | Negative |
| 13 | F | 21 | Negative |
| 14 | M | 19 | Negative |
| 15 | F | 20 | Negative |
| 16 | F | 21 | Negative |
| 17 | F | 20 | Negative |
| 18 | F | 22 | Negative |
| 19 | F | 22 | Negative |
| 20 | F | 32 | Negative |
| 21 | F | 23 | Negative |
| 22 | F | 20 | Negative |
| 23 | F | 20 | Negative |
| 24 | M | 21 | Negative |
| 25 | M | 20 | Negative |
| 26 | M | 21 | Negative |
| 27 | M | 22 | Negative |
| 28 | M | 21 | Negative |
| 29 | M | 25 | Negative |
| 30 | F | 21 | Negative |
| 31 | F | 22 | Negative |
| 32 | M | 28 | Negative |
| 33 | F | 21 | Negative |
| 34 | F | 22 | Negative |
| 35 | F | 18 | Negative |
| 36 | F | 18 | Negative |
| 37 | F | 22 | Negative |
| 38 | M | 23 | Negative |
| 39* | F | 21 | Positive |
| 40* | F | 19 | Negative |

\*Volunteers 39 and 40 have been excluded from the study due to poor compliance

